# Machine learning (decision tree analysis) identifies ecological selectivity patterns across the end-Permian mass extinction

**DOI:** 10.1101/2020.10.09.332999

**Authors:** William J. Foster, Georgy Ayzel, Terry T. Isson, Maria Mutti, Martin Aberhan

## Abstract

Decision tree algorithms are rarely utilized in paleontological research, and here we show that machine learning algorithms can be used to identify determinants of extinction as well as predict extinction risk. This application of decision tree algorithms is important because the ecological selectivity of mass extinctions can reveal critical information on organismic traits as key determinants of extinction and hence the causes of extinction. To understand which factors led to the mass extinction of life during an extreme global warming event, we quantified the ecological selectivity of marine extinctions in the well-studied South China region during the end-Permian mass extinction using the categorized gradient boosting algorithm. We find that extinction selectivity varies between different groups of organisms and that a synergy of multiple environmental stressors best explains the overall end-Permian extinction selectivity pattern. Extinction risk was greater for genera that were limited to deep-water habitats, had a stationary mode of life, possessed a siliceous skeleton or, less critically, had calcitic skeletons. These selective losses directly link the extinction to the environmental effects of rapid injections of carbon dioxide into the ocean-atmosphere system, specifically the combined effects of expanded oxygen minimum zones, rapid warming, and ocean acidification.

## 1 Introduction

Machine learning is one of the major subfields of artificial intelligence, classically defined as computational algorithms that have the ability to learn from data as to improve performance or make accurate predictions (Mohri et al., 2018). Furthermore, machine learning techniques can be sub-divided into supervised learning, unsupervised learning, or semi-supervised learning techniques. In this study we apply decision tree algorithms, which are some of the simplest, but powerful, supervised machine learning algorithms that can solve both regression and classification problems (Breiman et al., 1984). Decision tree algorithms offer immense potential to contribute to problem solving in geosciences (Karapatne et al., 2017) but have only been utilized in limited ways in paleontological research, such as fossil identification (Renaudie et al., 2018), text-mined data collection (Kopperud et al., 2019) and niche modelling (MacGuire and Stigall, 2009). Decision Tree algorithms can also be used to predict survival during mass mortality events. A key example is the Titanic dataset (www.kaggle.com/titanic), used to train data scientists in predicting survivorship based upon the metadata of passenger information (e.g., gender, wealth, age). Despite the potential of decision tree algorithms to predict extinction risk and its many applications across biology (e.g., Hanna and Cardilo, 2014), there have been few studies that have utilized this technique for paleobiology (Finnegan et al., 2012; Tietje and Rödel, 2018). Besides generating a model to predict survivorship, decision tree algorithms can also be used to rank ecological factors that are most important when making the predictive model and, thus, can reveal extinction selectivity patterns. Despite the potential of decision tree algorithms, these methods have only been utilized for exploring extinction selectivity during one mass extinction (i.e., the end-Ordovician mass extinction; Finnegan et al., 2012). To demonstrate the potential of decision tree algorithms in revealing extinction selectivity, we explore the ecological selectivity during the end-Permian mass extinction using the categorized gradient boosting decision tree algorithm.

The end-Permian mass extinction event is synchronous with one of the most extreme climate warming events of the Phanerozoic, which occurred approximately 252 million years ago (Burgess and Bowring, 2015). The timing of this extinction event coincides with the emplacement of the Siberian Traps large igneous province into carbon-rich sedimentary rocks, and the associated injection of large volumes of greenhouse gases into the atmosphere (Svensen et al., 2009). Based on our understanding of the impact of rapid and high magnitude carbon dioxide emissions, as well as both sedimentological and geochemical proxies from the Permian/Triassic boundary, it has been hypothesized that the causes of marine extinctions stem from ocean acidification (Payne et al., 2007), thermal stress (Joachimski et al., 2012), and deoxygenation (Wignall and Twitchett, 1996).

One way to explore the cause(s) of a mass extinction event is to investigate the ecological extinction selectivity. Several biological and ecological traits appear to have been selected against during the end-Permian mass extinction. Specifically, selectivity patterns of extinction have been recognized amongst physiology (Knoll et al., 2007; Clapham and Payne, 2011; Payne et al., 2016), mineralogy (Clapham and Payne, 2011), motility (Clapham, 2017), and tiering (Clapham and Payne, 2011). These selective patterns have led to suggestions that decreasing pH, hypercapnia and global ocean warming played a major role in the extinctions of marine organisms (Knoll et al., 2007; Clapham and Payne, 2011; Payne et al., 2016; Clapham, 2017). Furthermore, a biogeographic selectivity pattern has been related to the metabolic rate of marine organisms, suggesting that thermal stress in the tropics and deoxygenation towards the poles best explain the pattern of extinction (Penn et al., 2019). It is, however, unlikely that the end-Permian extinction selected equally strongly against all these different ecological attributes and some of these traits appear selected against because many of the traits are shared amongst the taxa that went extinct. Here, by applying a decision tree algorithm, the relative importance of the determinants of extinction can be resolved, which will highlight the most important selectivity patterns for the end-Permian event.

To investigate extinction selectivity patterns across the end-Permian mass extinction, we examined the ecological traits that were selected against to infer extinction mechanisms. The intensely studied and species-rich rock record of marine successions from South China provides an excellent opportunity to compare selectivity patterns between pre-extinction and the extinction interval while minimizing geographic, rock-record, and sampling biases. It is particularly important that South China covers a broad region with numerous sections and facies representing time intervals prior to the mass extinction that also occur in other sections afterwards, which allows for a consistency with both the type of marine environment and paleolatitude of the investigated Permian-Triassic successions. This limits the impact of a facies control on fossil occurrence patterns and any sequence stratigraphical overprints on the extinction pattern. In addition, since 63% of pre-extinction Changhisingian and 41% of post-extinction Griesbachian occurrences in the Paleobiology Database (paleobiodb.org) are derived from China, and because China is one of the only regions that records both a continuous and fossiliferous deposition across the Permian/Triassic boundary, this high volume of data for China biases our ‘global’ understanding of the end-Permian mass extinction (Fig. 1). This region, therefore, offers the best perspective on extinction dynamics for this event.

**Figure 1.**
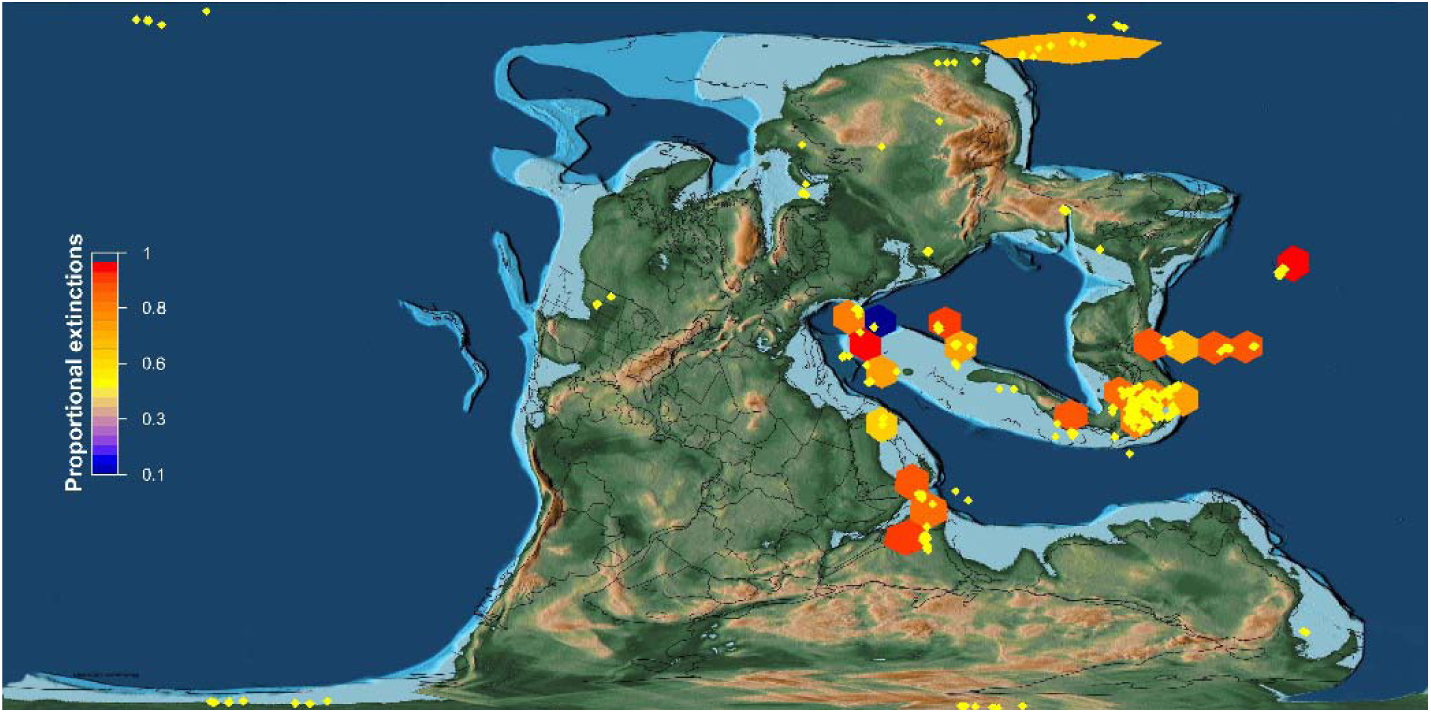
Proportional extinctions of marine genera during the end-Permian mass extinction event in different regions. Each hexagon cell represents an equal area and only cells that include both pre- and post-extinction data are included. The yellow diamonds represent pre-extinction sections with a fossil record. The plot was generated using data from the Paleobiology Database (paleobiodb.org) and plotted using the icosa package (Kocsis et al., 2018) in R overlain on a basemap from Scotese (Scotese, 2016).

## 2 Methods

We used a database of all marine invertebrate and calcareous algae genera recorded from the late Permian to the Middle Triassic in South China downloaded from the Paleobiology Database and supplemented with additional data from the literature (Fig. S1). The database includes 25683 occurrences of 1283 genera representing Brachiopoda, Foraminifera, Mollusca, Conodonta, Radiolaria, Arthropoda, Chlorophyta, Rhodophyta, Porifera, Cnidaria, Echinodermata, Bryozoa, and Annelida. Vetting of the data meant that undetermined genera, and informal taxa were excluded from the database. Latest synonymies and re-identifications were used where possible, and for mollusks were updated following MolluscaBase (molluscabase.org). Given that metadata information (e.g., depositional environment, section name, and formation name) were sometimes inconsistent among data sources even for the same fossiliferous bed in the same section, or because it was absent from the Paleobiology Database, the metadata were updated for each section using the most up-to-date literature for consistency. Occurrences of taxa from outside of the South China region that extend the stratigraphic ranges of genera into the Permian or into the Triassic were not included, to avoid introducing spatial variation and biases in the dataset. In some sections, in particular the Permian/Triassic boundary GSSP in Meishan, single beds have been subsequently divided into sub-beds which relate to different conodont biozones. Therefore, in older references that did not subdivide these beds the occurrence is considered to have occurred in all respective biozones. Where possible, using the original references, each occurrence was assigned to a conodont biozone (Fig. S1). All analyses were carried out at the genus-level, because of increased likelihood of misidentifications at the species-level, 29% of species occurrences in our database lack an identification, and the high proportion of singleton species. For all the analyses genera with single occurrences were omitted. Extinction rates were calculated for those genera that can be assigned to conodont biozones, that is 12889 occurrences of 634 genera. Extinction rates were calculated following Foote (2000): extinction rate = -log[*N*_*bt*_/(*N*_*bL*_ + *N*_*bt*_)], where *N*_*bL*_ is the number of taxa that cross the bottom boundary of a time bin only and *N*_*bt*_ is the number of taxa that cross both boundaries.

To investigate extinction selectivity, we characterized each genus according to ten ecological traits (Table 1) and one phylogenetic criterion (phylum) using the primary literature, extant relatives, and the Paleobiology Database referring to the ecological attribute during a taxon’s adult life stage (see also supplemental material). Genera that exhibited ecological traits that differed between species, e.g., differences in the strength of ornamentation, were split into different taxa to consider within-genus ecological variation. To analyze extinction selectivity, the data were binned into four time intervals due to the nature of the end-Permian mass extinction event (see results) and to avoid over-splitting the data: the Wuchiapingian, pre-extinction Changhsingian, the mass extinction interval (Changhsingian *C. yini* to the Griesbachian *H. parvus* Zone; as defined by Wang et al., 2014), and the Griesbachian post-extinction interval.

**Table 1.**
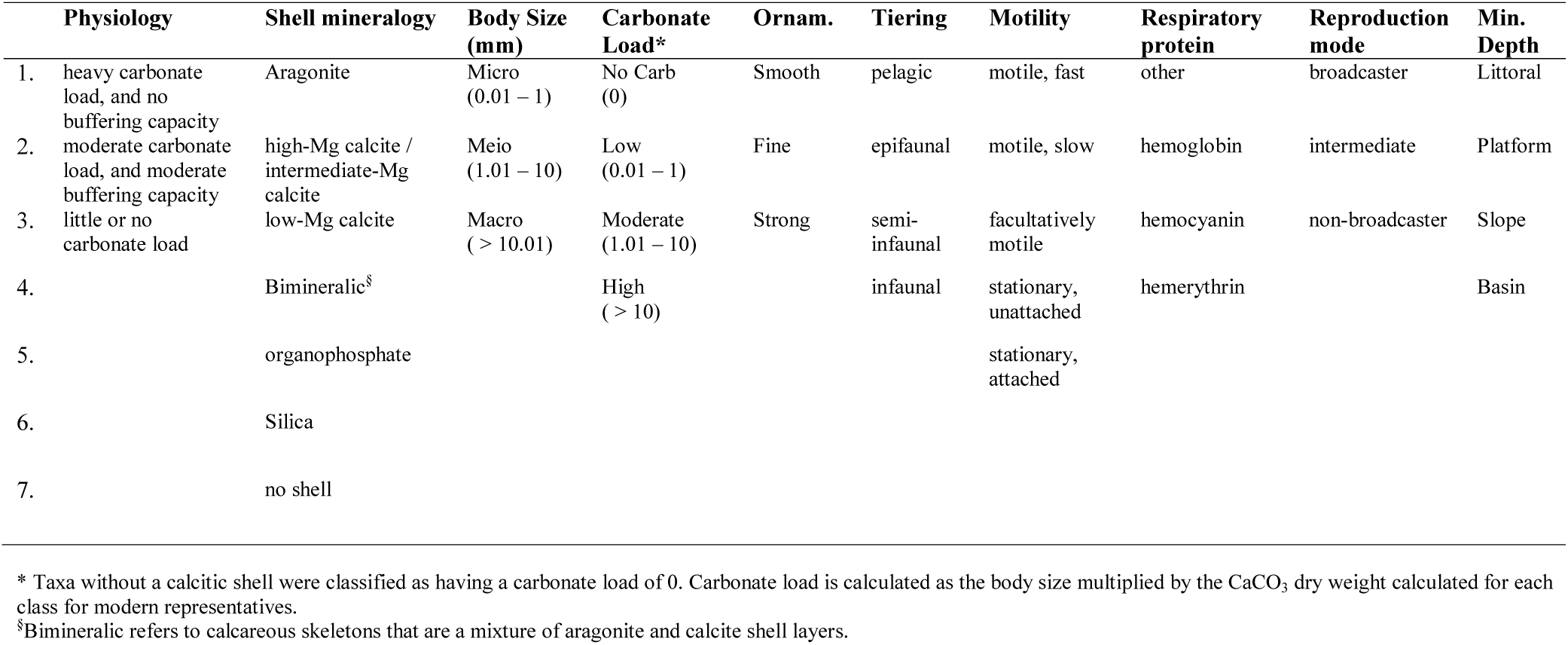
Ecological categories used in this study. Physiological buffering capacity after Knoll et al. (2007) and Payne et al. (2016). Tiering and motility after Bambach et al. (2007). Ornamentation after Aberhan et al. (2006). Reproduction mode after Bush and Bambach (2016). Descriptions for the remaining categories are given in the supplementary material. Physiology and shell mineralogy are ranked according to the expected sensitivity to ocean acidification following Knoll et al. (2007) and Ries (2011), respectively. Ornam. = ornamentation. Min. Depth = minimum depth.

To identify and rank the key drivers of extinction risk we applied the categorical gradient boosting decision trees (CGB-DT) method. CGB-DT is a powerful and high-performance machine learning technique with a better prediction performance in comparison with conventional linear models (Ayaru et al., 2015; Belyaeva et al., 2017). The core idea behind the CGB-DT technique is to combine simple decision tree models with gradient boosting (Friedman, 2001) in the way that the next decision tree is trying to minimize the error of the previous decision tree. In the present study, we used a CGB-DT model to predict extinction risk as a binary classification problem, i.e., whether a species went extinct (“1”) or not (“0”) based on the set of corresponding traits. To fit the CGB-DT model we used the open-sourced machine learning library for gradient boosting CatBoost (Dorogush et al., 2018; Prokhorenkova et al., 2018). We trained an individual CGB-DT model for each investigated time interval with default CatBoost hyperparameters, 5-fold cross-validation (i.e., splits, where data are split into a training and evaluation subset), and the log loss function as an optimization criterion. We estimated the obtained model performance with the area under the receiver operating characteristic (AUC) which is sensitive to type I and type II error. An AUC is a measure of discrimination between two distinct groups of species we try to classify as extinctions or survivors, thus, it is closely related to the Wilcoxon signed-rank test which determines whether two randomly dependent samples have the same distribution. An AUC was averaged over the splits we used for cross-validation. Additionally, we performed recursive feature elimination analysis (as proposed in Belyaeva et al., 2017) which confirmed the robustness of the proposed approach in ranking traits. As we expect correlations among our variables, prior to model building we checked the predictors for multicollinearity (Fig. S2). A correlation plot of the different predictors shows that a few predictors are correlated, e.g., body size and carbonate load during the extinction interval (Fig. S2). Correlation values are, however, low indicating that no undesirable levels of multicollinearity are present.

Machine learning algorithms are often described as a black box due to the lack of transparency associated with how algorithms make predictions. To demonstrate how the CGB-DT algorithm generate its predictions and to increase interpretability, we applied shapely additive explanations (SHAP) values to exemplify the output from the CGB-DT algorithm (Lundberg and Lee, 2016). First, we investigated the collective SHAP values for each variable, which highlights how much each predictor contributes to the target variable, and the individual SHAP values for each genus, which shows how the contribution of different features affects the prediction.

All the data and code regarding performed machine learning modeling and analysis are available on GitHub (https://github.com/hydrogo/mel). Each part of our research workflow can be reproduced interactively using Binder at https://mybinder.org/v2/gh/hydrogo/mel/master.

## 3 Results

### 3.1 Nature of the end-Permian mass extinction

Exploring the selectivity of the end-Permian mass extinction first requires the timing and the duration of the event to be determined. The mass extinction event has been described as a single event in the latest Permian (Jin et al., 2000), two separate events extending into the earliest Triassic (Song et al., 2013), or a single extended extinction interval spanning the Permian/Triassic boundary (Wang et al., 2014). Our analyses of extinction rates in South China show that those genera that can be correlated to conodont biozones have heightened extinction rates in the *Clarkina yini* Zone, in the *Hindeodus zhejiangensis* – *H. changxingensis* zones, and throughout most of the Induan (Figs. 2, S3), but a low extinction rate in the *C. meishanensis* Zone (Fig. 2). This low extinction rate in the *C. meishanensis* Zone also corresponds with a relatively low number of occurrences compared to surrounding zones, and the low extinction rate is likely a consequence of insufficient sampling. Given that the highest extinction rates occurred in the extinction interval as defined by Wang et al. (2014), we follow Wang et al. (2014) and interpret the extinction event as representing an interval spanning the P/Tr boundary, which also avoids over-splitting the data. This suggests that the extrinsic changes that occurred over this approx. 60 kyr interval (Burgess and Bowring, 2015) caused the end-Permian mass extinction. Consequently, to investigate extinction selectivity, the data were aggregated into four time intervals: the Wuchiapingian, pre-extinction Changhsingian, the mass extinction interval (Changhsingian *C. yini* to the Griesbachian *H. parvus* Zone; Wang et al., 2014), and the Griesbachian post-extinction interval.

**Figure 2.**
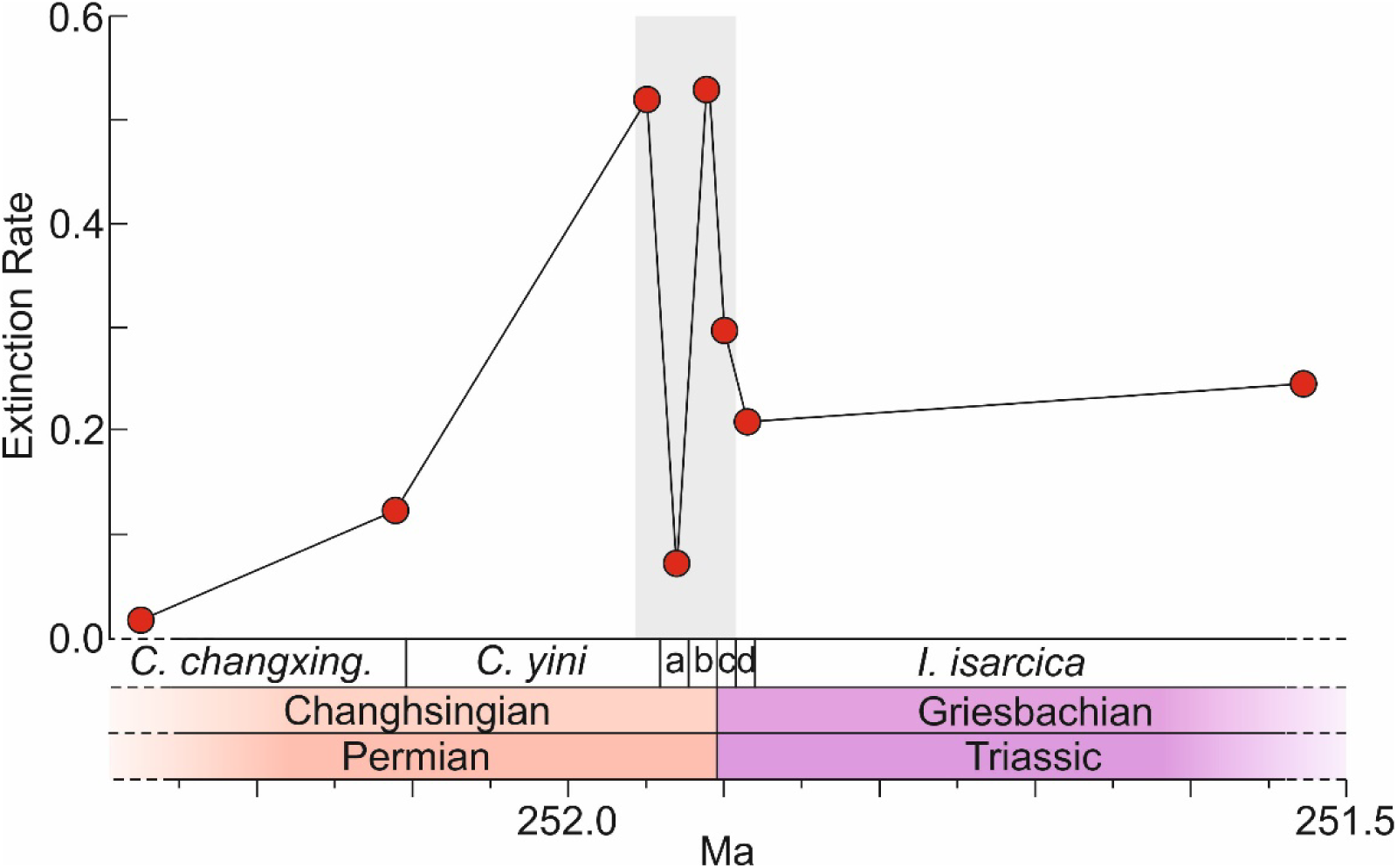
High-resolution extinction rates of marine genera across the Permian/Triassic boundary in South China. The extinction interval is highlighted in gray after Wang et al. (2014). Radiometric ages after Burgess and Bowring (2015). Conodont zones after Yuan et al. (2014); a = *Clarkina meishanensis*, b = *Hindeodus zhejiangensis* – *H. changxingensis*, c = *H. parvus*, and d = *Isarcicella staeschei. C. changxing*. = *C. changxingensis*. Extinction rates for the full studied interval are shown in Fig. S3.

### 3.2 Extinction selectivity

The area under the curve-receiver operating characteristics curve (AUC) (Fig. 3) visualizes the CGB-DT model performance. This curve plots the true positive rate against the false positive rate. An AUC of 0.5 and indicates the performance of a random classification that has no utility (Fig. 3). It is unlikely that a decision tree algorithm will give an AUC value of 1 which represents a perfect classification, instead an AUC > 0.7 is typically considered representative of a good model (Mohri et al., 2018). The AUC for each time interval (Fig. 3), show an average of 0.68, 0.76, 0.75, and 0.74 for the Wuchiapingian, pre-extinction Changhsingian, mass extinction interval, and Griesbachian post-extinction time interval, respectively. In particular, the curves for the pre-extinction Changhsingian and the mass extinction interval show that the CGB-DT model is a good classification model for interpreting extinction selectivity.

**Figure 3.**
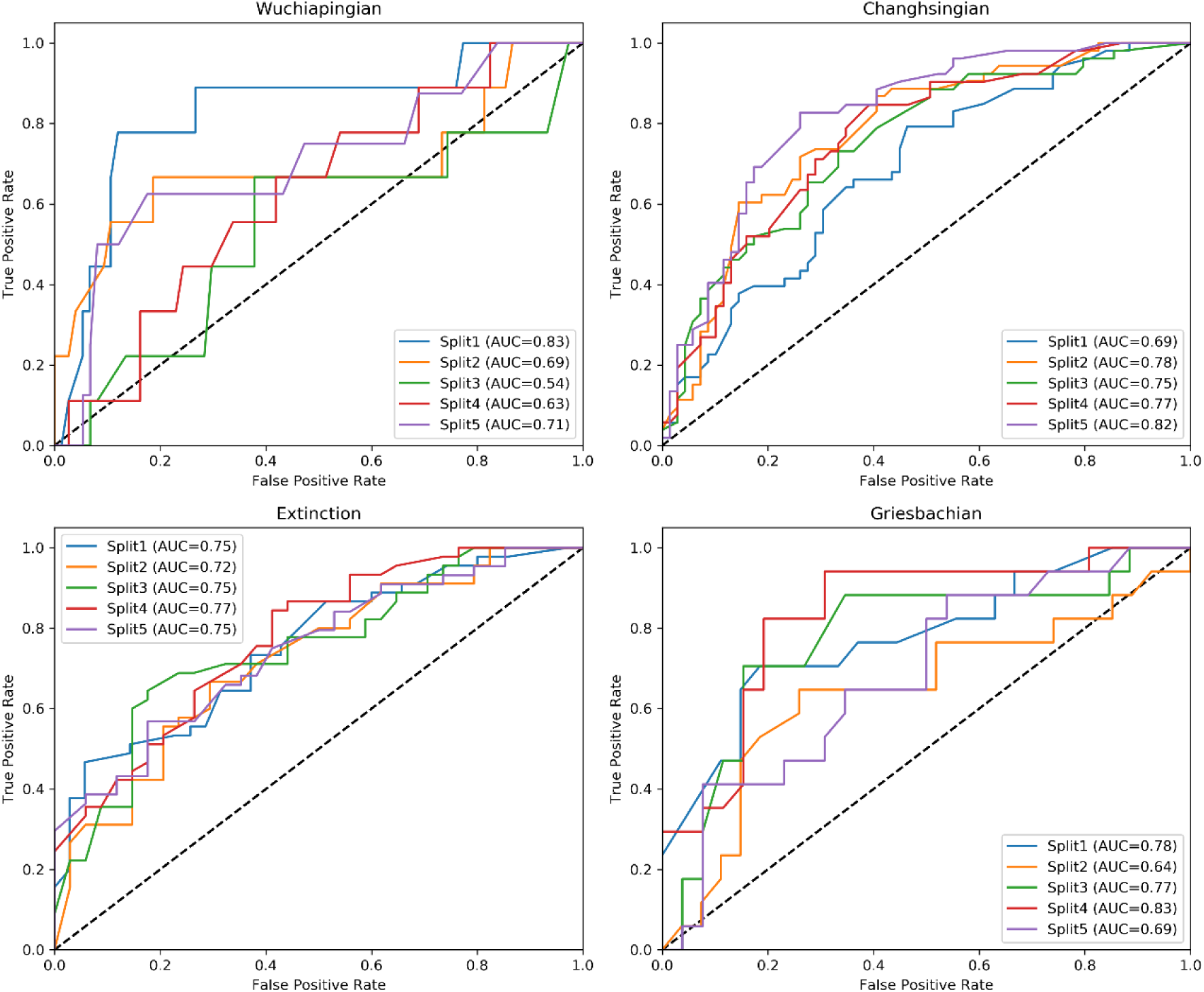
Area under the curve-receiver operating characteristics curves (AUC) for the splits used for cross-validation in each time interval. The black dashed line represents an AUC of 0.5 which indicates a model with a random classification.

CGB-DT reveals that the relative importance of the ecological variables associated with extinction varies between intervals (Fig. 4A; S4). The difference in the relative importance of factors between the pre-extinction interval and the extinction interval indicates that the end-Permian extinction was not simply triggered by an intensification of pre-extinction pressures, a characteristic shared with other mass extinction events (Finnegan et al., 2012; Dunhill et al., 2018). The relative importance of different factors for the extinction interval highlights that skeletal mineralogy, motility, and physiology are the best predictors of extinction risk. These were also better predictors of extinction than phylum, showing that these ecological attributes are more significant in predicting extinction than phylogenetic membership. Habitat depth was also an important predictor of extinction prior to the extinction interval (Fig. 4), suggesting that one of the main drivers of ecological selectivity prior to the extinction interval was still important during the extinction interval.

**Figure 4.**
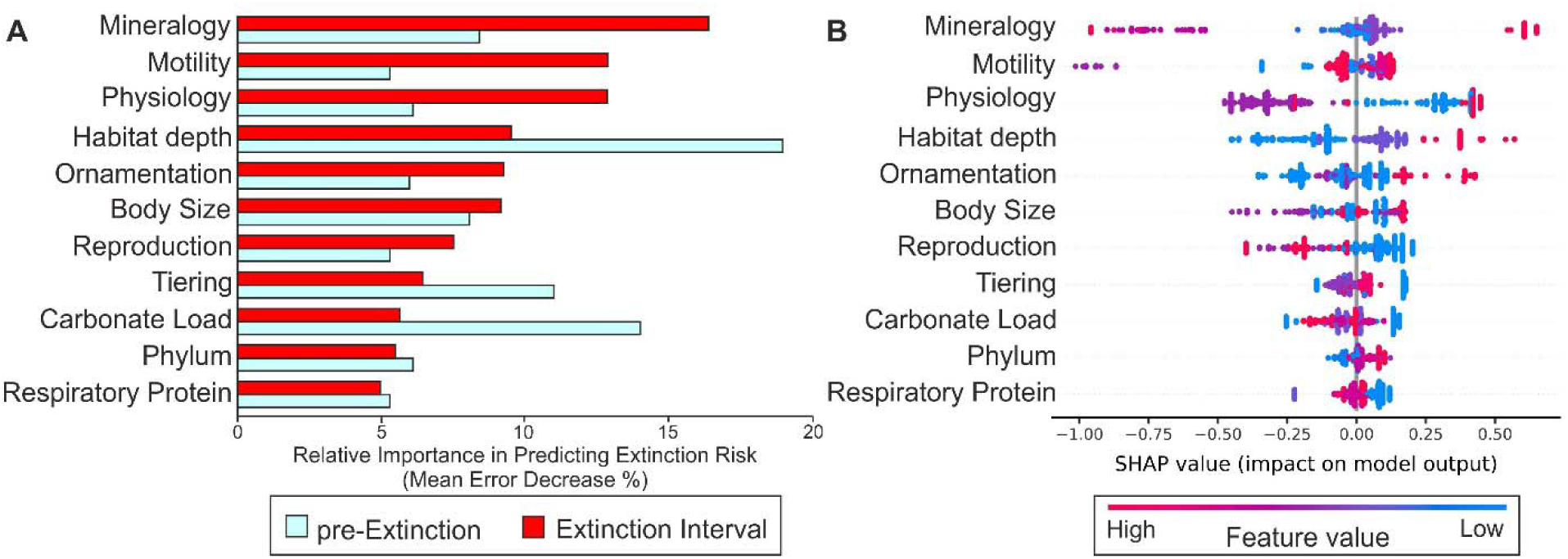
Extinction selectivity based of ecological attributes of marine genera from South China. A) The relative importance of each ecological variable. B) SHAP summary plot showing how the difference values of each ecological attribute effect the model predictions for the extinction interval. The horizontal location of the values shows if a data point from the training dataset is associated with a higher or lower prediction. The vertical position corresponds to the relative importance of each ecological attribute. The SHAP summary plot for split 3 is shown.

The end-Permian extinction was highly selective according to an organisms mineralogy, with the SHAP summary plot indicating that the CGB-DT would predict a lower extinction risk for genera with no shell or an organophosphatic shell and a higher extinction risk for genera with a siliceous shell (Fig. 4B). This can also be seen in the SHAP values attributes for individual genera where mineralogy for these mineralogical groups has one the greatest importance in model predictions (Fig. 5A-C). The selectivity of the other mineralogical groups is less clear (Fig. 6B). Aragonite, high-Mg calcite, and low-Mg calcite genera are more likely to go extinct than the remaining mineralogical groups, but this only has a small influence on the CGB-DT (Fig. 4B; Fig. 5D).

**Figure 5.**
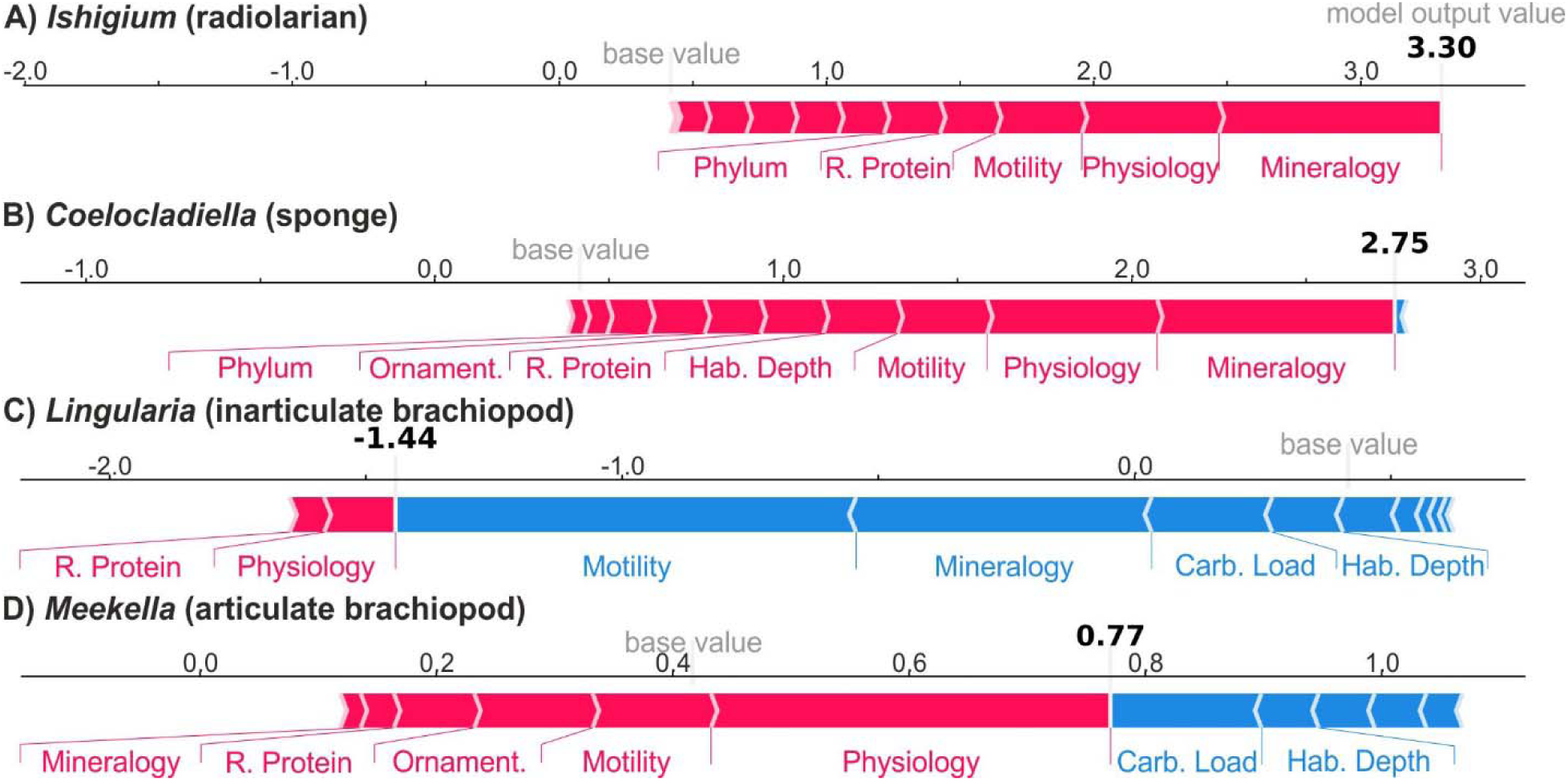
Relative SHAP value attributes showing how the different ecological variables for four example genera change in the model prediction for the extinction interval. The base value is the prediction if no ecological variables are considered. Model output value is the prediction considering the ecological variables for the investigated genus, with positive values indicating extinction and negative values indicating survival. Features that push the prediction higher (to the right), i.e., more likely to go extinct, are shown in red, and the opposite is shown in blue. Carb. Load = carbonate load, Hab. Depth = habitat depth, R. Protein = respiratory protein.

**Figure 6.**
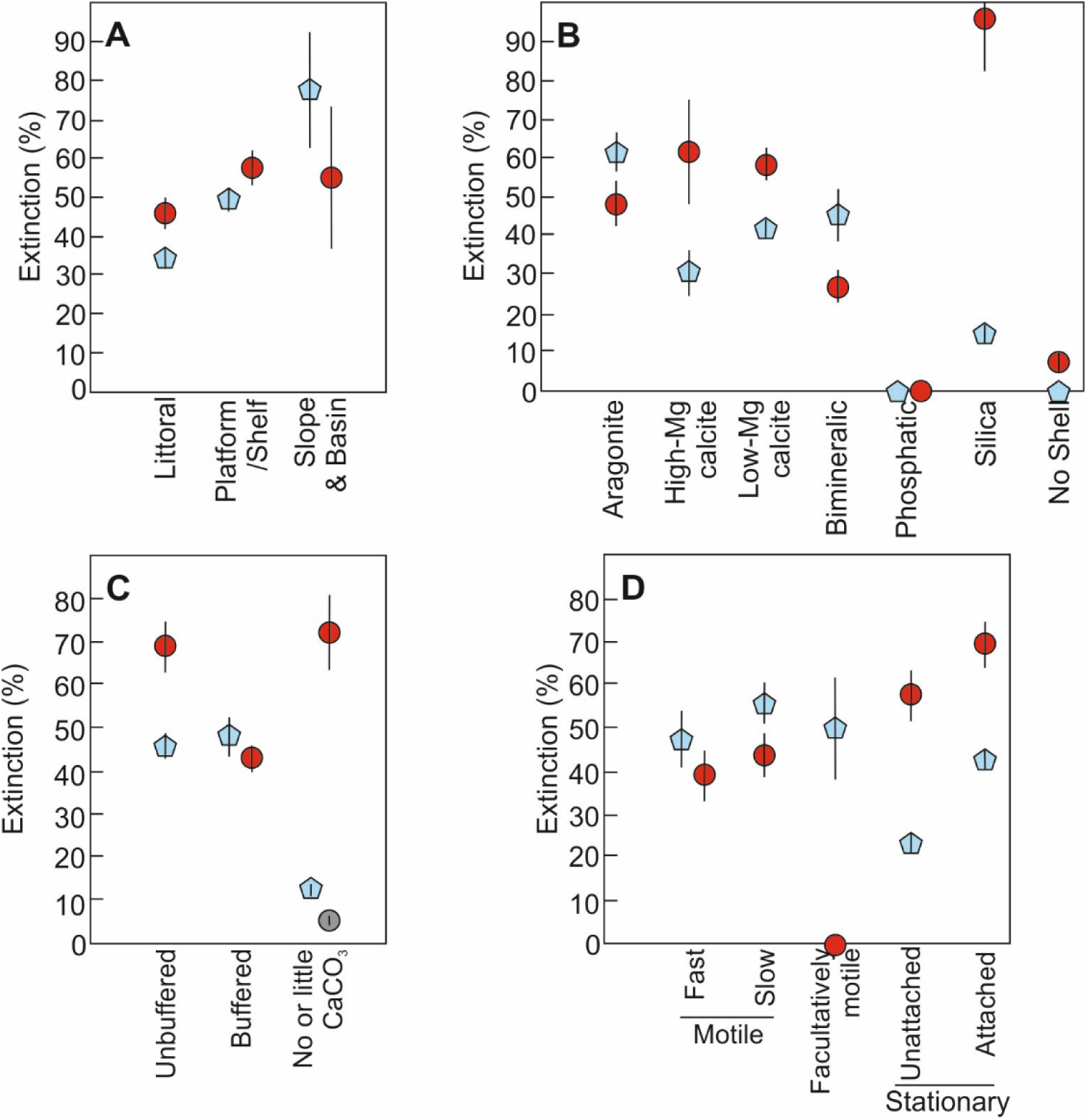
Percent extinction of marine genera with different habitat preferences and organismic traits during the pre-extinction Changhsingian (blue pentagon) and the extinction interval (red circles) in South China. A, minimum water depth, B, skeletal mineralogy, C, physiology, the grey circle indicates extinction risk of taxa with no or little carbonate load after siliceous genera were removed, and D, motility.

The differences in percentage extinctions among different habitat depths show that prior to the mass extinction there is a clear shallow-deep water selectivity gradient with those genera confined to basinal settings being more likely to go extinct than those that ranged into shallow settings on the platform or close to the shoreline (Fig. 6A). Because of these extinctions, genus richness in basinal settings was low during the extinction interval and the selectivity pattern appears to disappear. The SHAP summary plot (Fig. 4B) shows, however, that this selectivity pattern still exists during the extinction interval and that the reduced importance in the CGB-DT model is because the generic richness of the deeper settings is lower than in the Changhsingian.

Physiology and motility are equally important for the CGB-DT model during the extinction interval (Fig. 4A). When considering physiology, it is clear that genera with a heavy carbonate load and no buffering capacity where highly selected against, whereas animals with a moderate carbonate load and a moderate buffering capacity were less likely to go extinct (Figs. 4B; 6C). Genera with little or no carbonate load, however, show a mixed signal because this group includes siliceous taxa that were highly selected against. This means that for non-siliceous genera, e.g., the articulate brachiopod *Meekella* (Fig. 5D), physiology is a good predictor of extinction risk, whereas for siliceous taxa mineralogy is a better predictor of extinction risk (Fig. 5A-B), consistent with previous studies (Knoll et al., 2007; Clapham and Payne, 2011; Payne et al., 2016). When looking at the extinction magnitude between the different types of motility, genera that are motile are less likely to have gone extinct than stationary genera (Fig. 6D). The CGB-DT also shows that for facultatively motile taxa, motility is the most important factor for predicting a low extinction risk (Fig. 4B).

## 4 Discussion

The extinction selectivity according to an organism’s habitat depth in both the pre-extinction and extinction interval was high, with a clear shallow-deep water selectivity gradient (Figs. 4B; 6A). The selective loss of benthic genera restricted to deeper water coincides with elevated extinction rates during the Changhsingian (prior to the mass extinction interval). As indicated by redox proxies from South China (Song et al., 2012), this is tied to the expansion of oxygen minimum zones and the concomitant reduction of habitable areas in basinal settings (He et al., 2015). The loss of these genera from basinal settings is not considered a consequence of a sea-level change, sampling-bias, or rock record-bias because post-extinction basinal facies in South China have also been intensely studied but do not yield body fossils. Shoaling of the carbonate compensation depth (CCD), which is the depth at which carbonate is undersaturated leading to dissolution of marine shells, may also explain the selective loss of genera in basinal settings, as increases in dissolved CO_2_ will lead to calcium carbonate undersaturation. Notably, some basinal successions became devoid of carbonate in the upper Changhsingian (He et al., 2005), which has been inferred as deposition below the CCD (Isozaki, 1997). Understanding the processes that govern the shoaling and stability of the CCD during the Paleozoic is, however, speculative because changes in the position of the CCD in modern oceans involves important pelagic calcifiers, i.e, planktic foraminifera and coccolithophores, that did not become an important component of marine ecosystems until the mid-Mesozoic. During the extinction interval, shallow platform settings (below fair-weather wave-base) were also exposed to transient anoxic conditions (Song et al., 2012), which may explain the increased extinction risk in this setting (Figs. 4B; 6A). The selective loss of genera restricted to deep-water habitats suggests that the expansion of the oxygen minimum zone in basinal settings was one of the main drivers of extinction.

The end-Permian extinction was highly selective against siliceous taxa compared to calcareous genera (Figs. 4B; 6B). A drop in the diversity and abundance of both radiolarians and silicisponges led to a crash in biosiliceous ooze production resulting in an equatorial chert gap during the Early Triassic (Racki, 1999). One factor that could have led to the demise of effective silica factories during the end-Permian extinction is rapidly rising ocean temperatures that would have increased silica dissolution rates and altered the silica saturation state (Racki, 1999; Beauchamp and Baud, 2002). Temperature stress is also expected to produce selective patterns of extinction by preferentially extinguishing genera that already lived near their upper thermal limits (Song et al., 2014). Based on what is known on the thermal limits of modern radiolarians (Song et al., 2014) the temperatures that developed during the extinction interval in South China (Joachimski et al., 2012) likely made equatorial surface waters uninhabitable. Further support that high temperatures caused this selective signal is that radiolarians, silicisponges, and chert deposits that do occur after the mass extinction event are only known from thermal refugia, i.e., high paleolatitudes or deeper water settings (Godbold et al., 2017; Takemura et al., 2002), and that in shallow marine settings the major metabolic stress in the tropics was thermal stress (Penn et al., 2019).

The preferential extinction of siliceous genera does not negate ocean acidification, and instead highlights that rapid increase in CO_2_ into the ocean-atmosphere system has numerous complex consequences on ecosystems. Experimental studies investigating the impact of ocean acidification on the biological calcification of marine animals have shown that species which precipitate aragonite and high-Mg calcite are more vulnerable than those that precipitate low-Mg calcite (Ries, 2011). Even though our results show that genera that precipitate siliceous skeletons were more susceptible to extinction than any other mineralogical group, aragonite, high-Mg calcite and low-Mg calcite genera were also significantly selected against when compared to genera that possess no shell (Fig. 6B). Despite this selectivity pattern that is consistent with an ocean acidification scenario, the slightly higher extinction likelihood of genera with low-Mg calcite skeletons over aragonite skeletons during the extinction interval is inconsistent with modern studies (e.g., Ries et al., 2009; Ries, 2011) suggesting that ocean acidification was not a selective pressure during the extinction interval.

It is not only a genus’ CaCO_3_ polymorph that dictates whether it is vulnerable to ocean acidification. For instance, an organisms’ ability to regulate pH and carbonate chemistry at the site of calcification, the degree to which the organisms’ biomineral is protected by an organic coating, and shell microstructure are characteristics that have also been noted (Ries et al., 2009; Ries, 2011; Garbelli et al., 2017). Even though these factors are important it is not yet possible to include these in a detailed deep-time extinction selectivity analysis as this information is largely unknown, even for extant organisms. Physiological classifications to include these factors have been performed, but only at the class level (Knoll et al., 2007). These studies found a selectivity pattern consistent with an ocean acidification event. Using the same physiological categories, our study found a selective pattern that is inconsistent with ocean acidification and previous studies, i.e., genera with little or no carbonate load were more strongly selected against than ‘buffered’ genera with calcareous skeletons (Fig. 6C), owing to the selective loss of siliceous genera. When siliceous taxa are removed from the study, however, the extinction pattern is consistent with an ocean acidification scenario (Fig. 6C). Our study also shows that physiology was one of the most important ecological factors for predicting extinction in non-siliceous taxa (e.g., *Meekella*, Fig. 5D). Notably, the few studies that have been able to compare some of these additional factors within phyla have found a selectivity pattern consistent with ocean acidification, e.g., differences in the shell microstructure of Strophomenata and Rhynchonellata brachiopods (Garbelli et al., 2017).

The remaining variable that had a strong effect on extinction risk is motility. Genera that are motile are less likely to have gone extinct than stationary genera (Fig. 6D), consistent with previous observations (Foster and Twitchett, 2014; Clapham, 2017). The resilience of motile genera to rapid climate warming is likely due to motile genera typically having a greater aerobic scope, inherently high extracellular *p*CO_2_, and higher maximum thermal tolerance limits (Clapham, 2017). The selective loss of stationary genera did not exist prior to the mass extinction which suggests that conditions responsible for this selective pattern developed rapidly during the extinction interval. The preferential extinction of stationary genera compared to motile taxa cannot be used to infer individual extinction mechanisms, as the physiological adaptations of motile taxa are advantageous under high-temperature, low-oxygen, and low-pH conditions.

Recently the biogeographical and physiological selectivity of the end-Permian mass extinction has been interpreted to have been a consequence of a combination of high temperatures and widespread marine anoxia (Song et al., 2014; Penn et al., 2019). Our CGB-DT results, however, show that ecological selectivity during the end-Permian extinction varies among different groups of organisms and that a synergy of multiple environmental stressors best explains the overall end-Permian extinction selectivity patterns. Given that the selectivity patterns observed for the extinction interval in this study are interpreted as a consequence of expanded oxygen minimum zones, high sea-surface temperatures, and ocean acidification, this selective pattern corroborates the results of previous studies (i.e., Knoll et al., 2007; Clapham and Payne, 2011) that suggest multiple factors synergistically drove the end-Permian mass extinction event and that this “Deadly Trio of carbon dioxide” is responsible for the ecological selectivity signal at the end-Permian extinction.

## 5 Conclusions

Decision tree algorithms have a great potential to reveal selectivity patterns during both past and projected extinction events. Applying the categorical gradient boosting algorithm to a database of marine invertebrates that span the Permian/Triassic boundary in South China, our analysis reveals that extinction risk was greater for genera that were limited to deep-water habitats, had a stationary mode of life, possessed a siliceous skeleton or, less critically, had calcitic skeletons. Furthermore, these selectivity patterns did not exist prior to the extinction suggesting that extinction drivers changed between the pre-extinction interval and the extinction interval. We linked these selective losses to the synergistic effects of rapid climate change associated with the end-Permian mass extinction, i.e., expanded oxygen minimum zones, rapid ocean warming, and ocean acidification.

## Acknowledgments

This project was funded by Geo.X grant SO_087_GeoX and is associated with the DFG Research Unit TERSANE (FOR 2332: Temperature-related stressors as a unifying principle in ancient extinctions). The authors also thank the numerous authors of the original studies that provide the source data on which this study is based, and the many data enterers of the Paleobiology Database for the provision of fossil occurrence data. We also thank Alexander Dunhill, Bethany Allen, Wolfgang Kiessling, Richard Twitchett, Matthew Clapham, Jonathan Payne, and an anonymous reviewer for their constructive comments on this research and manuscript.

## Supporting Information

**Figure S1.**
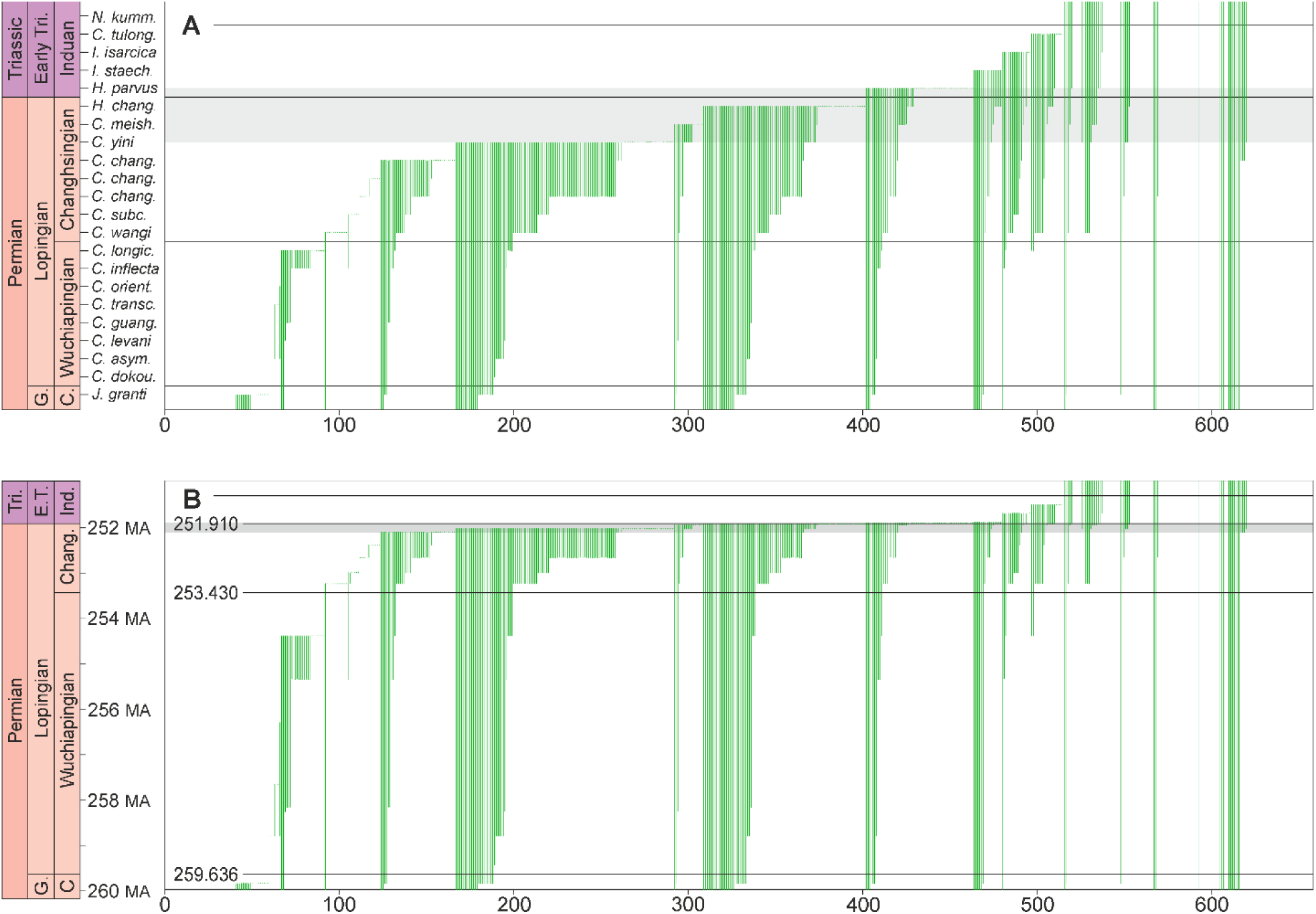
Stratigraphic ranges of fossil marine invertebrate genera used in this study. Genus numbers are shown on the x axis and are organized by the last appearance date (LAD). The extinction interval is highlighted in grey. Substage and stage boundaries are shown by horizontal black lines. A) Genus ranges scaled to conodont zones with each conodont zone given an equal thickness. G. = Guadalupian, Early Tri. = Early Triassic, C. = Capitanian. Conodont zones after Yuan et al. (2014), Jin et al. (2003) and Zhang et al. (2007): *C. dokou*. = *Clarkina dukouensis, C. asym*. = *C. asymmetrica, C. guang. = C. guangyuanensis, C. transc*. = *C. transcaucasia, C. orient*. = *C. orientalis, C. longic*. = *C. longicuspida, C. subc*. = *C. subcarinata, C. chang*. = *C. changxingensis, C. meish*. = *C. meishanensis, H. chang*. = *Hindeodus changxingensis* – *C. zhejiangensis, I. staech*. = *Isarcicella staeschei, C. tulong*. = *C. tulongensis* – *C. planata, N. kumm*. = *Neogondolella kummeli* – *N. krystyni*. B) Genus ranges scaled to radiometric dates. Radiometric dates after Burgess et al. (2014) and Yuan et al. (2014). Tri = Triassic, E.T. = Early Triassic, Chang. = Changhsingian. MA = million years before present.

### Extended Materials and Methods

The body size of each species was based on the reported sizes, or taken from figured specimens, and the maximum body size for the species with the most occurrences within a genus was taken to represent the genus’ body size. The geometric mean of the genus body size was then calculated following Jablonski (1996). Because most genera that span the Permian/Triassic boundary are thought to have adapted their body size in response to environmental changes (Twitchett, 2007), and because interpopulation variance is not captured by single measurements for each genus, body size was categorized into three semi-quantitative groups (Table 1). The carbonate load was calculated by multiplying the body size of each genus that precipitates a carbonate skeleton by the percentage of the ash-free dry mass to the total mass for each faunal group based on modern representatives. Genera that do not possess a carbonate skeleton, e.g., radiolarians, were categorized as having no carbonate load. Ornamentation is a factor that is commonly related to predation pressure, whereby strong ornamentation enhances resistance against shell-breaking (Aberhan et al., 2006). In this study, however, we include ornamentation as an ecological attribute that is also related to carbonate load, whereby genera with strongly ornamented calcareous shells are expected to have greater carbonate loads than smooth genera. To investigate if genera that were restricted to deeper water habitats were selected against, we determined each genus’ shallowest habitat depth. Shallowest habitat depth was categorized as littoral, platform/shelf, or slope and basin. Pelagic taxa were not included in Fig. 6. Respiratory proteins are variable in structure, oxygen affinity, and concentration among invertebrate groups (Herreid II, 1980). Because of the different responses of proteins to changes in pH, temperature, and salinity it is expected that invertebrates with certain pigments, such as hemerythrin, will be less impacted by hypoxia than invertebrates that do not possess this pigment as an oxygen carrier. This selectivity pattern has been suggested to explain why lingulid brachiopods flourished following the end-Permian mass extinction (Peng et al., 2007; Posenato et al., 2014). Variances in the respiratory protein are known between invertebrate groups, but the variance within classes, such as bivalves, is less well-known.

**Figure S2.**
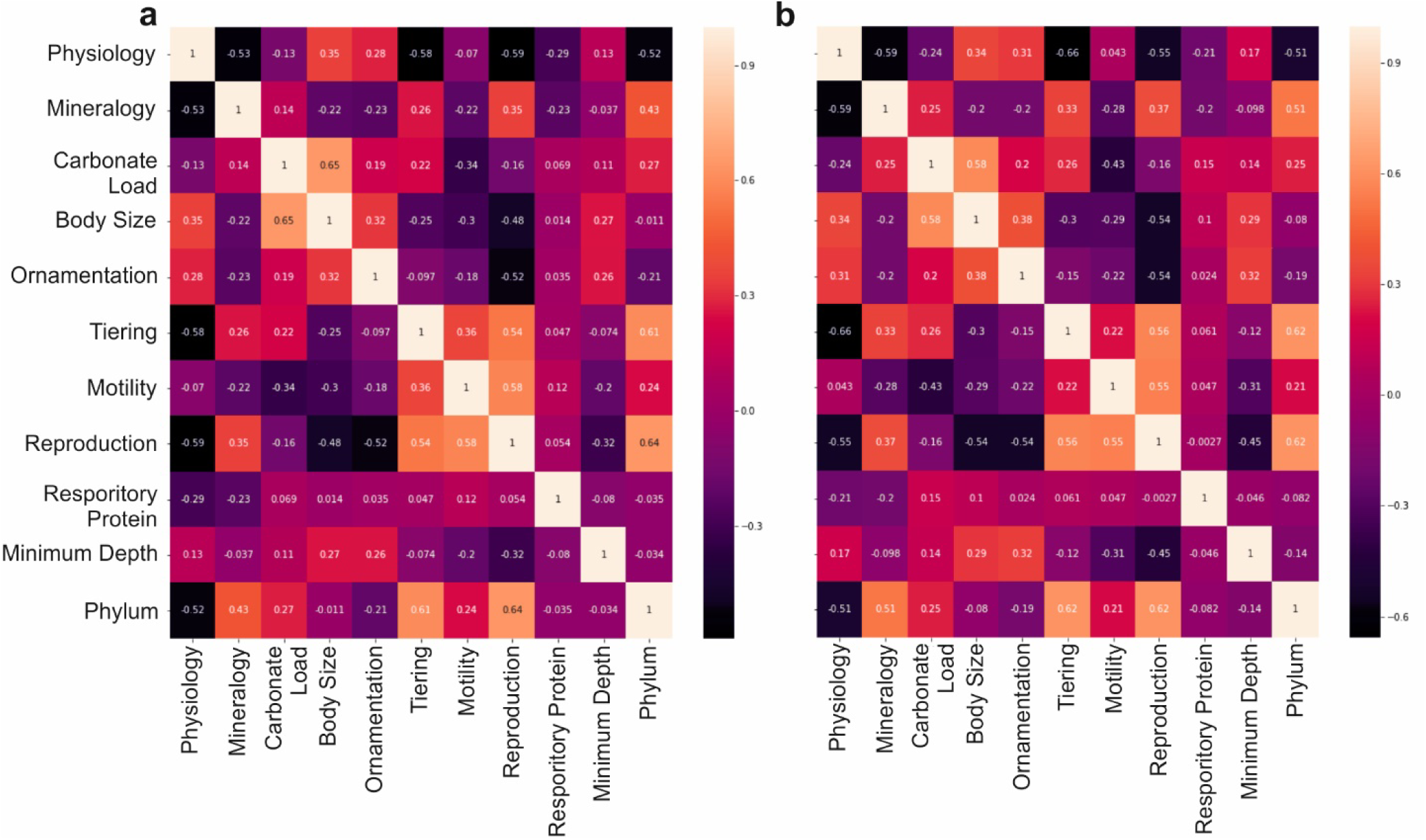
Correlation matrix of the investigated ecological attributes during (a) pre-extinction Changhsingian, and (b) the extinction interval.

**Figure S3.**
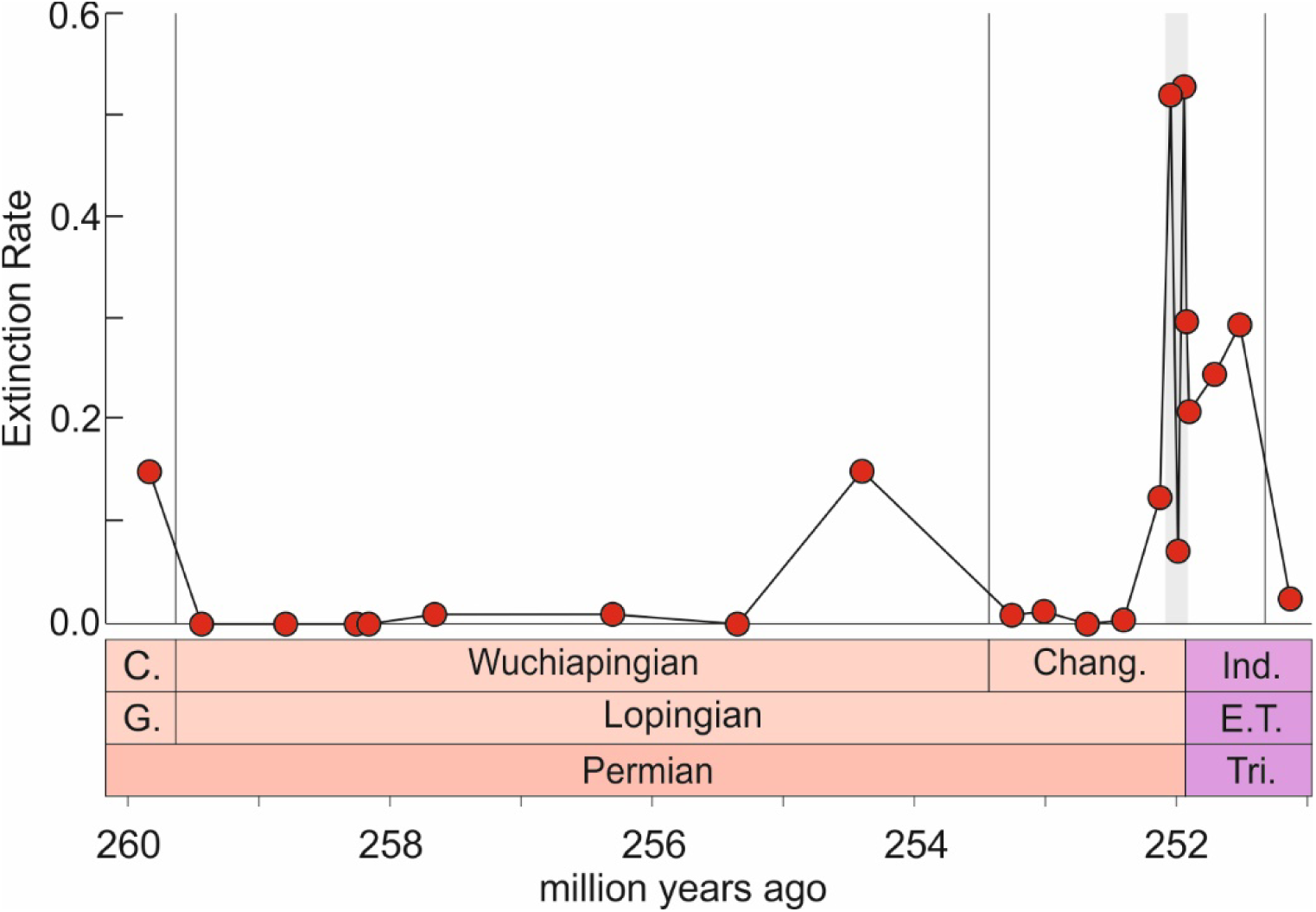
High-resolution regional extinction rates of marine genera across the studied interval in South China. Extinction rates are calculated after Foote (2000). The extinction interval is highlighted in grey after Wang et al. (2014). Radiometric ages after Burgess et al. (2014) and Yuan et al. (2014). Conodont zones after Yuan et al. (2014) See Figure 1 to zoom in on extinction rates across the Permian-Triassic boundary interval. C. = Capitanian, G. = Guadalupian. E.T. = Early Triassic, Ind. = Induan, Tri. = Triassic, Chang. = Changhsingian.

**Figure S4.**
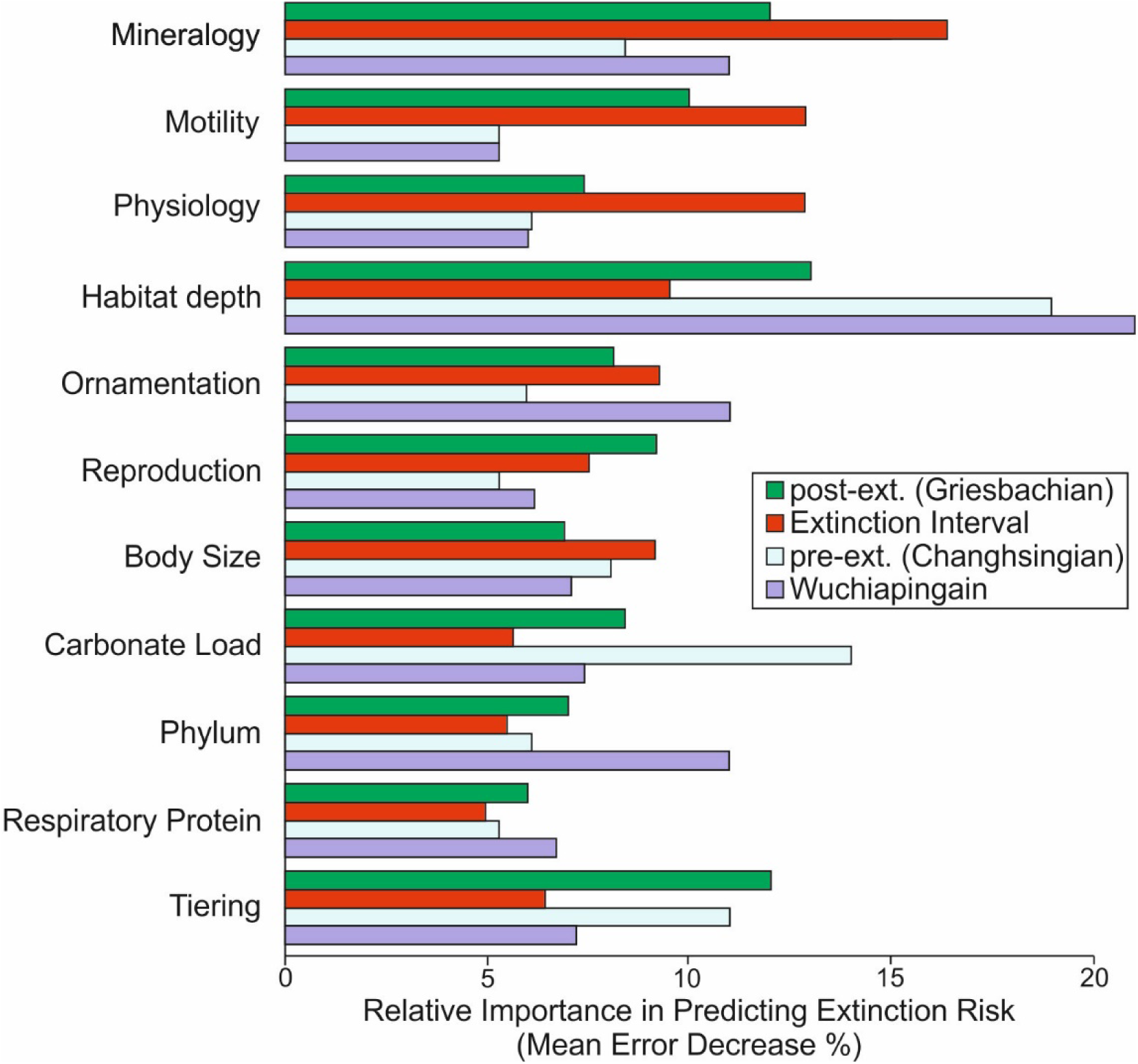
Relative importance of ten ecological traits and one phylogenetic attribute of marine genera for predicting extinction risk across the end-Permian mass extinction using high performance categorical gradient boosting on decision trees.

**Figure S5.**
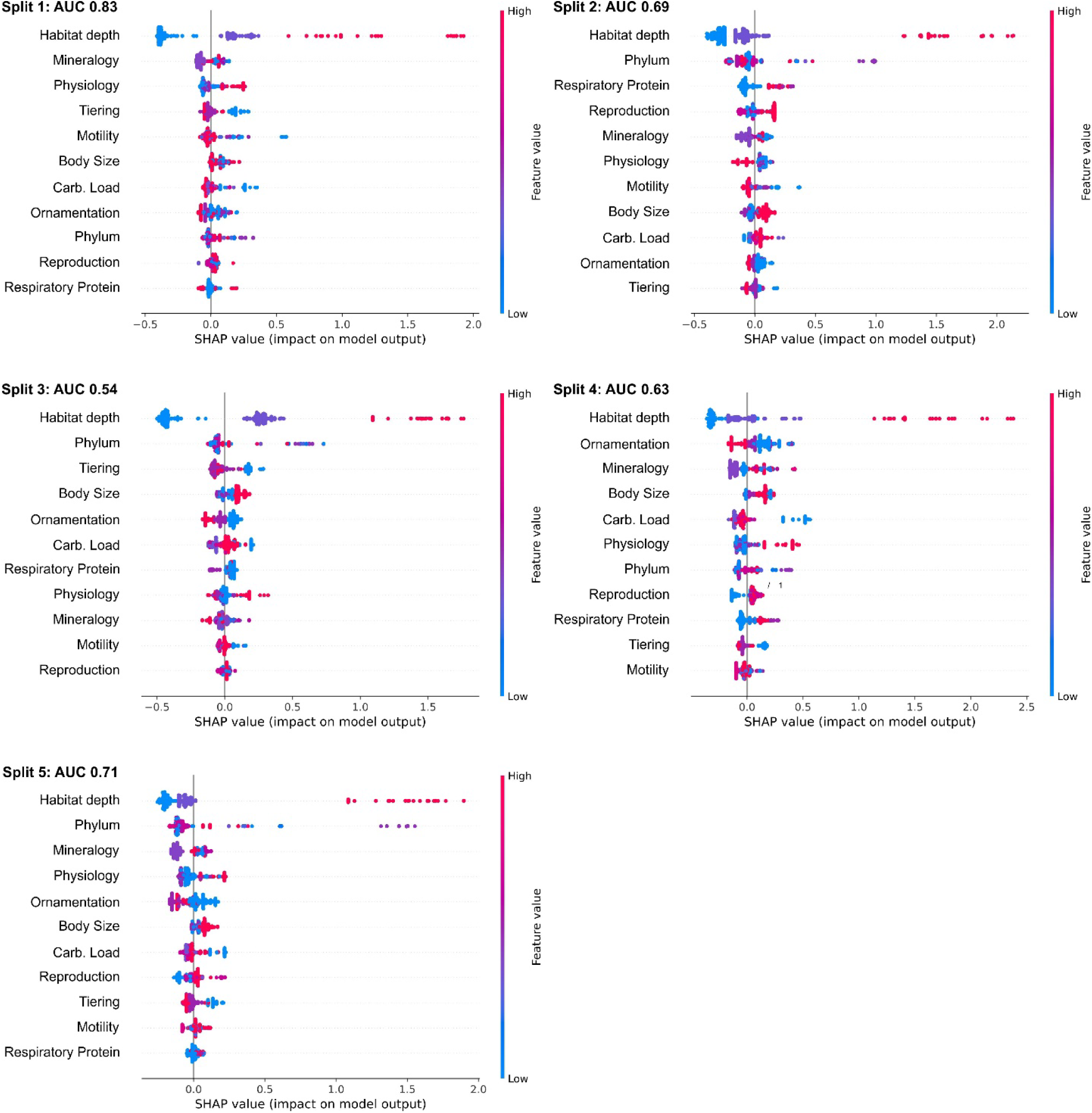
SHAP summary plots showing the relative importance of ecological attributes and how the different values for each ecological category affects the model predictions for the Wuchiapingian. The horizontal location of the values show if a data point from the training dataset is associated with a higher or lower prediction.

**Figure S6.**
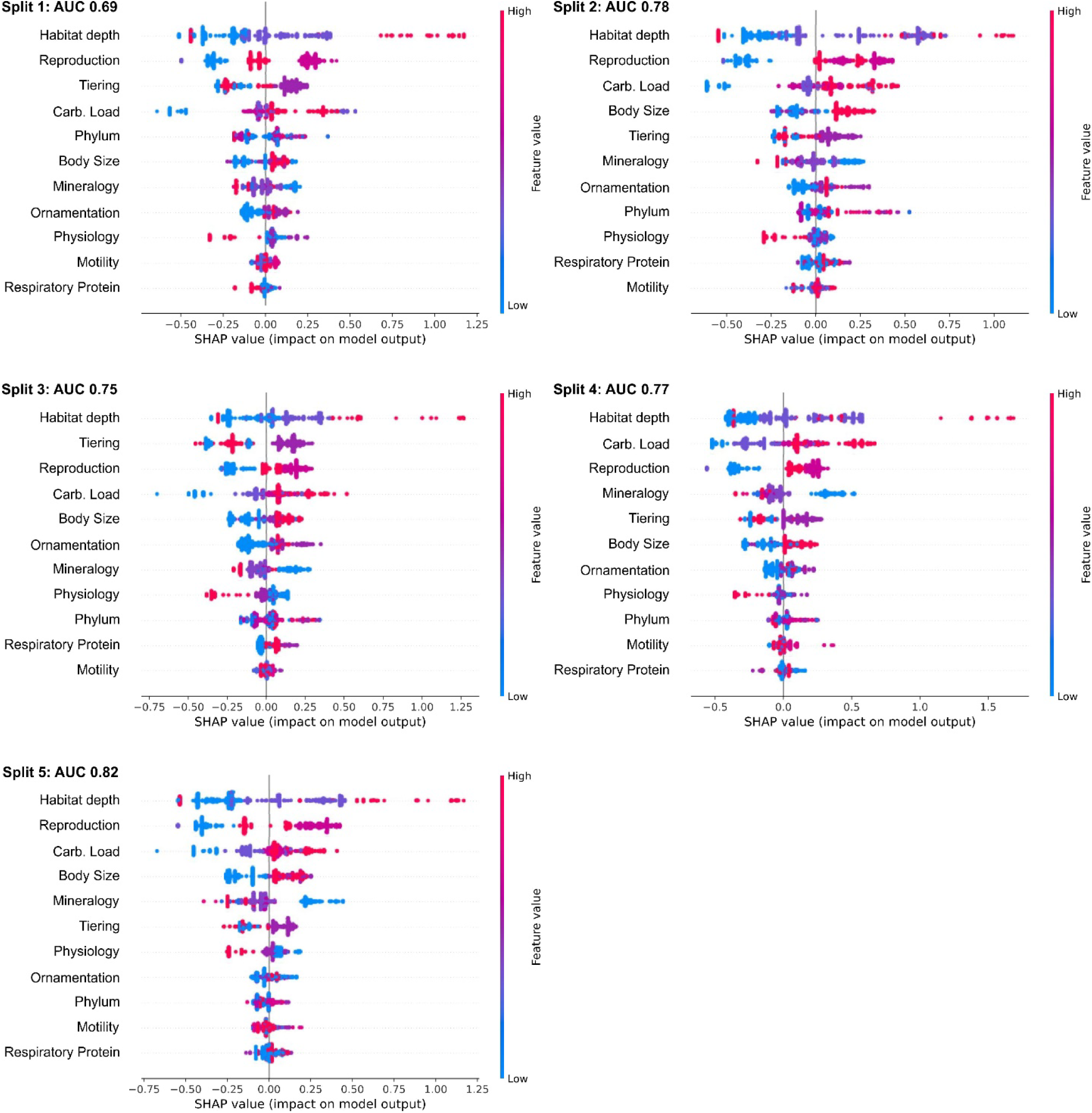
SHAP summary plots showing the relative importance of ecological attributes and how the different values for each ecological category affects the model predictions for the Changhsingian. The horizontal location of the values show if a data point from the training dataset is associated with a higher or lower prediction.

**Figure S7.**
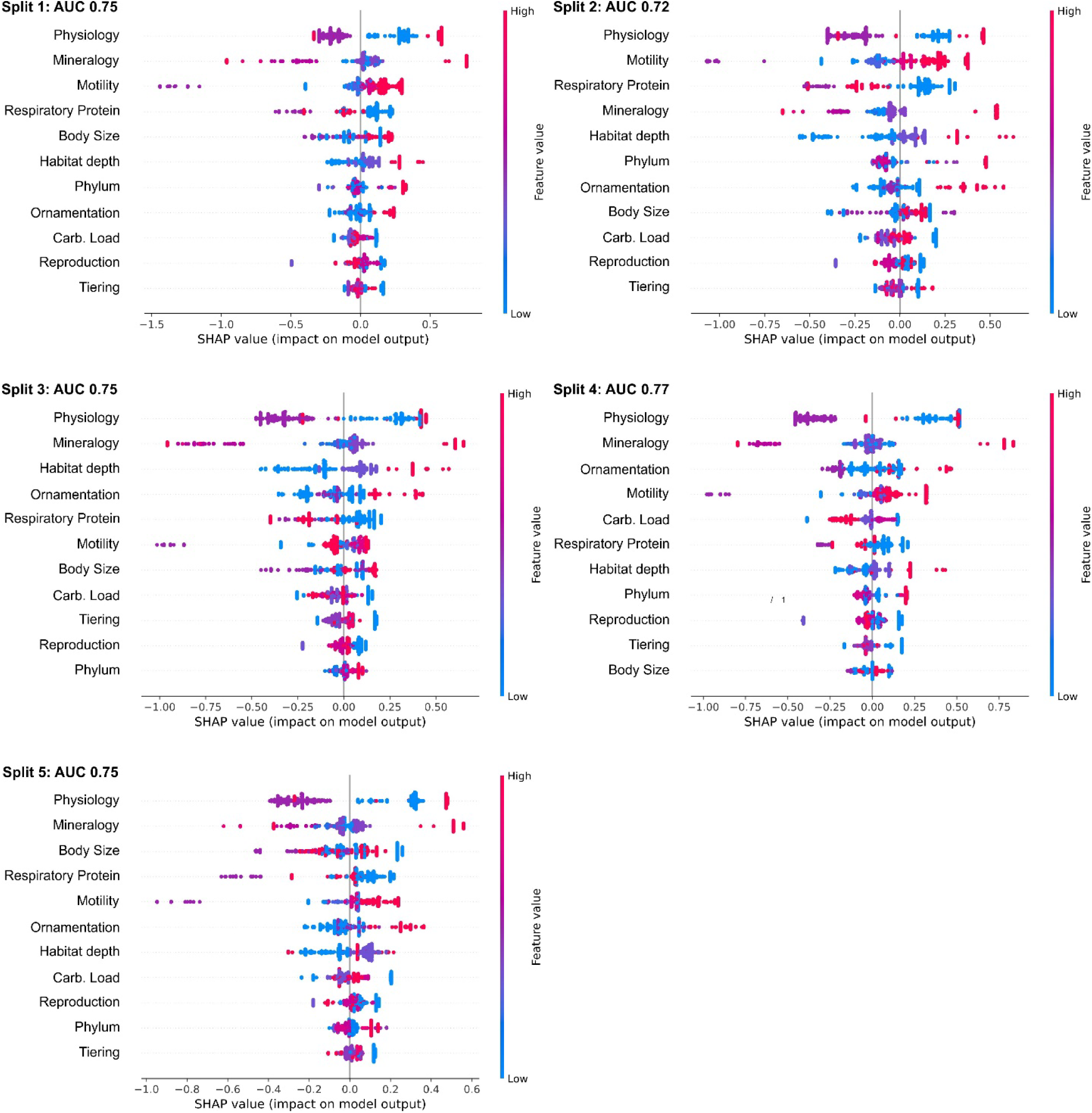
SHAP summary plots showing the relative importance of ecological attributes and how the different values for each ecological category affects the model predictions for the Extinction Interval. The horizontal location of the values show if a data point from the training dataset is associated with a higher or lower prediction.

**Figure S8.**
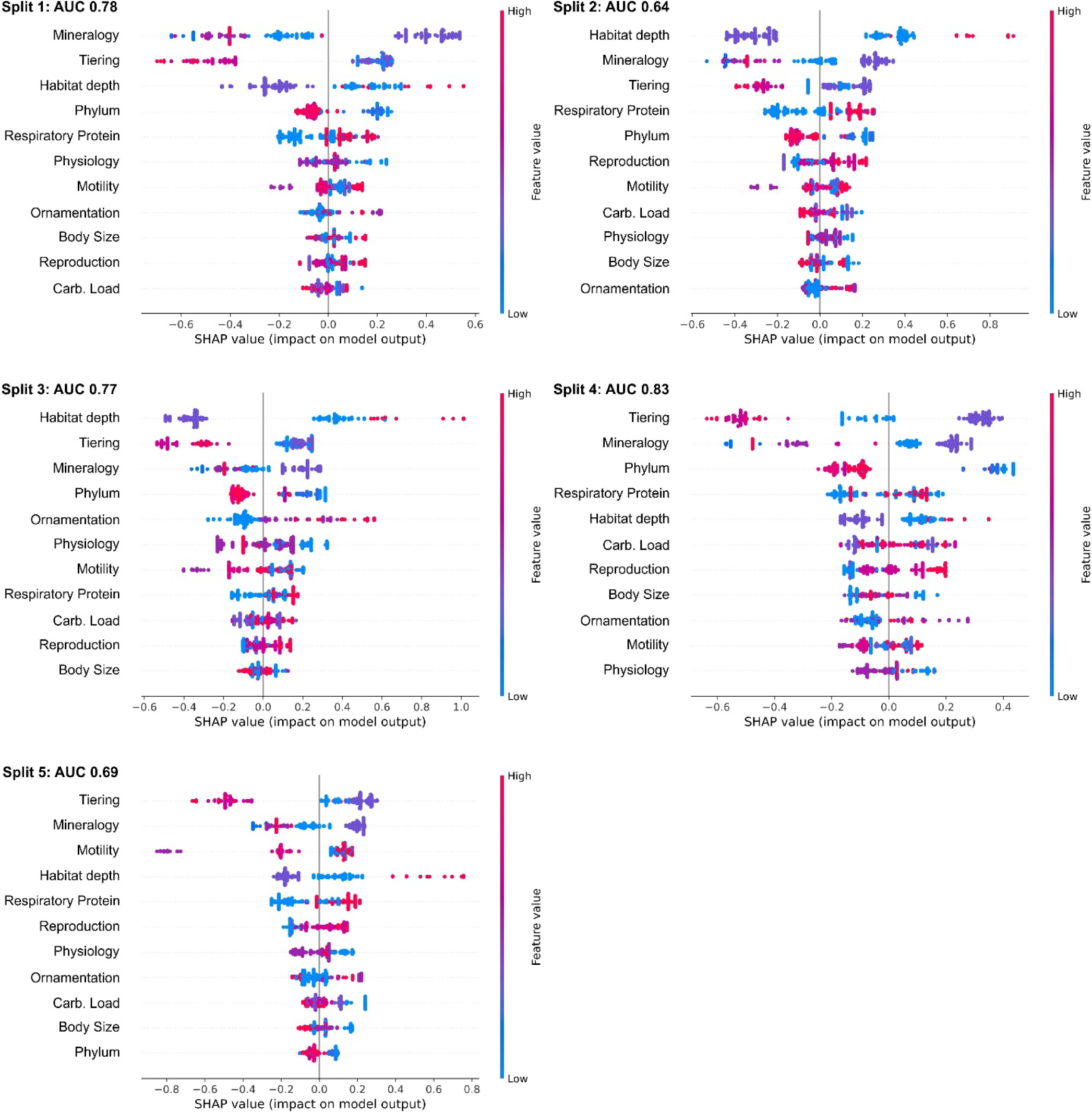
SHAP summary plots showing the relative importance of ecological attributes and how the different values for each ecological category affects the model predictions for the Griesbachian. The horizontal location of the values show if a data point from the training dataset is associated with a higher or lower prediction.

